# BRD9-containing non-canonical BAF complexes safeguard cell identity and prevent reprogramming

**DOI:** 10.1101/2021.05.27.445940

**Authors:** Kenan Sevinç, Gülben Gürhan Sevinç, Ayşe Derya Cavga, Martin Philpott, Simge Kelekçi, Hazal Can, Adam P. Cribbs, Enes Sefa Ayar, Dilşad H. Arabacı, James E. Dunford, Ata B. Demir, Logan H. Sigua, Jun Qi, Udo Oppermann, Tamer T. Onder

## Abstract

Epigenetic reprogramming requires extensive remodeling of chromatin landscapes to silence cell-type specific gene expression programs. ATP-dependent chromatin-remodeling complexes are important regulators of chromatin structure and gene expression; however, the role of Bromodomain-containing protein 9 (BRD9) and the associated ncBAF (non-canonical BRG1-associated factors) complex in reprogramming remains unknown. Here, we show that genetic suppression of BRD9 as well as ncBAF complex subunit GLTSCR1, but not the closely related BRD7, increase the efficiency by which induced pluripotent stem cells (iPSCs) can be generated from human somatic cells. Chemical inhibition and acute degradation of BRD9 phenocopied this effect. Interestingly, we find that BRD9 is dispensable for establishment and maintenance of human pluripotency but required for mesendodermal lineage commitment during differentiation. Mechanistically, BRD9 inhibition downregulates somatic cell type-specific genes and decreases chromatin accessibility at somatic enhancers. Collectively, these results establish BRD9 as an important safeguarding factor for somatic cell identity whose inhibition lowers chromatin-based barriers to reprogramming.

## Introduction

Expression of transcription factors (TFs) such as Oct4, Sox2, Klf4, c-Myc (OSKM) can erase somatic cell identity and reprogram the cells to a pluripotent state (*1, 2*). In doing so, reprogramming factors reset the entire epigenetic landscape established throughout development (*3*). The varying and low efficiencies of this process point to the presence of intrinsic somatic barriers to cell fate conversions. Several chromatin factors such as DOT1L methyltransferase (*4*), histone chaperone CAF-1 (*5*), BET family proteins (*6*), RNA Pol II regulator RPAP1 (*7*), SUMO modification (*8*), chromatin regulator FACT (*9*) and CBP/EP300 bromodomains (*10*) have emerged as potent barriers to reprogramming and act mainly by safeguarding pre-existing gene expression programs. Inhibition of these factors greatly facilitate reprogramming of a wide range of cell types (*11*). Discovery of additional safeguarding factors will likely yield important insights into chromatin-based mechanisms that maintain cell identity and restrict cell plasticity.

ATP-dependent chromatin remodeling complexes evict, exchange and space nucleosomes driven by the hydrolysis of ATP (*12*). Chromatin remodelers can facilitate transcriptional activation or repression based on the genomic location they bind to and additional chromatin factors they recruit (*13*–*15*). Among these, NuRD, INO80 and SWI/SNF complexes have been shown to modulate reprogramming in a variety of contexts (*16*–*19*). For example, overexpression of BAF complex subunits Smarca4 and Smarcc1 enhances murine somatic cell reprogramming by facilitating binding of Oct4 to its gene targets (*18*). In contrast, BAF complex subunits, Smarca2 and Smarcc2, have shown to be barriers in this context through upregulation of Stat3 and its target genes (*20*). Suppressing these somatic BAF subunits have been shown to activate the pluripotency circuit (*20*). These studies point to regulatory roles for different ATP-dependent chromatin remodeling complexes in various reprogramming frameworks.

Non-canonical BAF (ncBAF) complex is a recently identified SWI/SNF complex that lacks SMARCE, SMARCB, ARID1 and DPF but includes specific subunits such as BRD9 and GLTSCR1/L (*21*–*25*). BRD9 binds to enhancer regions in a cell type-specific manner and inhibition of its bromodomain leads to apoptosis in acute myeloid leukemia cells (*26*). Similarly, BRD9 inhibition leads to decreased cell proliferation, G1-arrest and apoptosis in rhabdoid tumor cells (*24*). In mouse embryonic stem cells (mESCs), ncBAF has been shown to co-localize with key regulators of naive pluripotency and BRD9 bromodomain inhibitors abolish naïve pluripotency by disrupting ncBAF’s recruitment to chromatin (*27*). These studies suggest that BRD9 is important for regulating cell identity and survival. However, the role of BRD9 and the ncBAF complex in somatic cell reprogramming remains unknown. In this study, we addressed this question using a combination of chemical and genetic tools in somatic cells and revealed an important role for BRD9 in safeguarding cell identity in the context of human reprogramming.

## Results

### Genetic suppression of ncBAF-specific subunits increases reprogramming efficiency

To investigate the role of ncBAF complex in reprogramming, we employed two genetic loss-of-function approaches. First, knockdown of *BRD9* using 2 independent shRNAs increased reprogramming efficiency 2-fold (fig. S1, A and B). On the other hand, suppression of *BRD9* paralog and PBAF-specific subunit, *BRD7*, had no effect on reprogramming even though it was efficiently knocked-down (fig. S1, A and C). In the second approach, we utilized sgRNAs and Cas9 to knock-out *BRD9* and then test the reprogramming efficiency of the resulting cells. Consistent with the shRNA data, knock-out of *BRD9*, but not *BRD7*, boosted reprograming efficiency up to 3-fold compared to control sgRNA expression (Fig. 1, A, B and C). Based on these results, we hypothesized that among the various BAF complexes in somatic cells, BRD9-containing ncBAF complex is a specific barrier for reprogramming. To test this notion, additional ncBAF specific subunits Glioma tumor suppressor candidate region gene 1 (GLTSCR1) or its paralog GLTSCR1-like (GLTSCR1L) were targeted by sgRNAs in human fibroblasts (*21*). Knocking out *GLTSCR1* increased reprogramming efficiency up to 4-fold for 3 sgRNAs out of 4 tested compared to control sgRNA expression (Fig. 1D). Targeting *GLTSCR1* paralog, *GLTSCR1L*, had no effect on reprogramming even though T7 endonuclease assay confirmed indel formation at this locus (Fig. 1D and fig. S1D). These results show that ncBAF complex members act as barriers to human somatic cell reprogramming.

**Figure 1.**
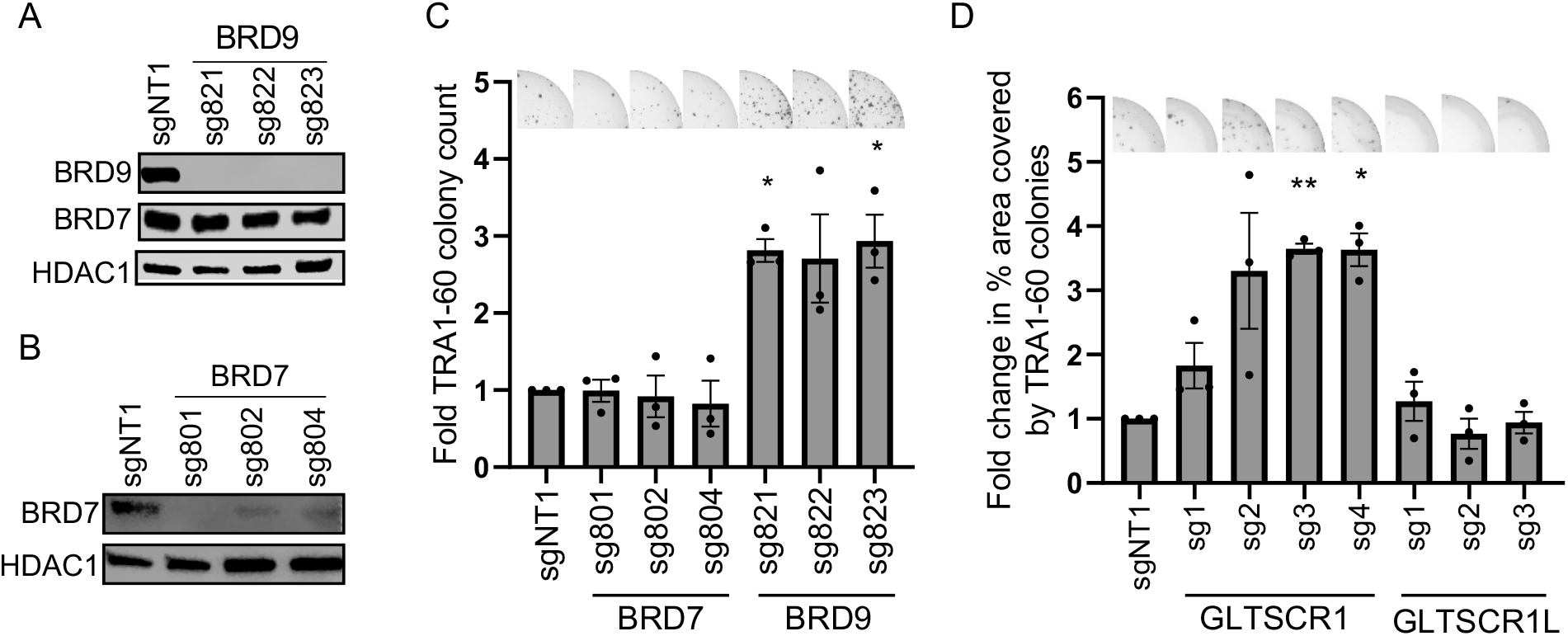
Genetic suppression of ncBAF-specific subunits increases reprogramming efficiency. **(A)** Western blots for BRD9 and BRD7 in fibroblasts expressing control (sgNT1) and BRD9-targeting gRNAs. HDAC1 serves as loading control. **(B)** Western blots for BRD7 in fibroblasts expressing control (sgNT1) and BRD7-targeting gRNAs. HDAC1 serves as loading control. **(C)** Fold change in the number of TRA-1-60-positive colonies for indicated sgRNA-expressing cells compared to control sgNT1-expressing cells. Representative well images are shown above the graph. Bar graphs show the mean and error bars represent standard error of mean. N=3, biological replicates. P-values are calculated by one sample t-test for mu=1. * denotes p<0.05, exact p-values from left to right are: 0.9593, 0.7910, 0.6173, 0.0065, 0.0968, 0.0302. **(D)** Fold change in percent area covered by TRA-1-60-positive colonies generated from fibroblasts expressing *GLTSCR1* or *GLTSCR1L* sgRNAs compared to control sgNT1. Representative well images are above relative bars. Bar graphs show the mean and error bars represent standard error of mean. N=3, three biological replicates. P-values are calculated by one sample t-test for mu=1. * denotes p<0.05, ** denotes p<0.005, exact p-values from left to right are: 0.1442, 0.1248, 0.0009, 0.0092, 0.4638, 0.4252, 0.7585.

### Bromodomain inhibition and degradation of BRD9 facilitate reprogramming

To confirm the role of BRD9 in somatic cell reprogramming, we next employed 3 structurally different inhibitors LP99, BI-7273 and I-BRD9 all of which selectively target the bromodomain of BRD9 and block its interaction with acetylated lysines (*28*–*30*) (Fig. 2A). All three BRD9 bromodomain inhibitors significantly increased reprogramming efficiency of human fibroblasts to iPSCs up to two-fold at concentrations of 1 and 3 μM (Fig. 2B). Importantly, I-BRD9 treatment had an additive effect with the inhibition of DOT1L, a potent reprogramming enhancer that we had previously identified (*4*). Combined inhibition of BRD9 and DOT1L led to a remarkable 5-fold increase in the number of iPSCs generated from human fibroblasts (fig. S2A). Emerging iPSC colonies in both control and BRD9 bromodomain inhibitor-treated cultures exhibited canonical characteristics of pluripotency such as retroviral transgene silencing and expression of pluripotency-specific markers OCT4, SSEA4 and NANOG (fig. S2, B and C). When injected into immunodeficient mice, iPSCs derived from I-BRD9-treated fibroblasts were able to form teratomas containing differentiated cells from all three germ layers (fig. S2D). These results show that inhibition of BRD9 bromodomain enhances human somatic cell reprogramming.

**Figure 2.**
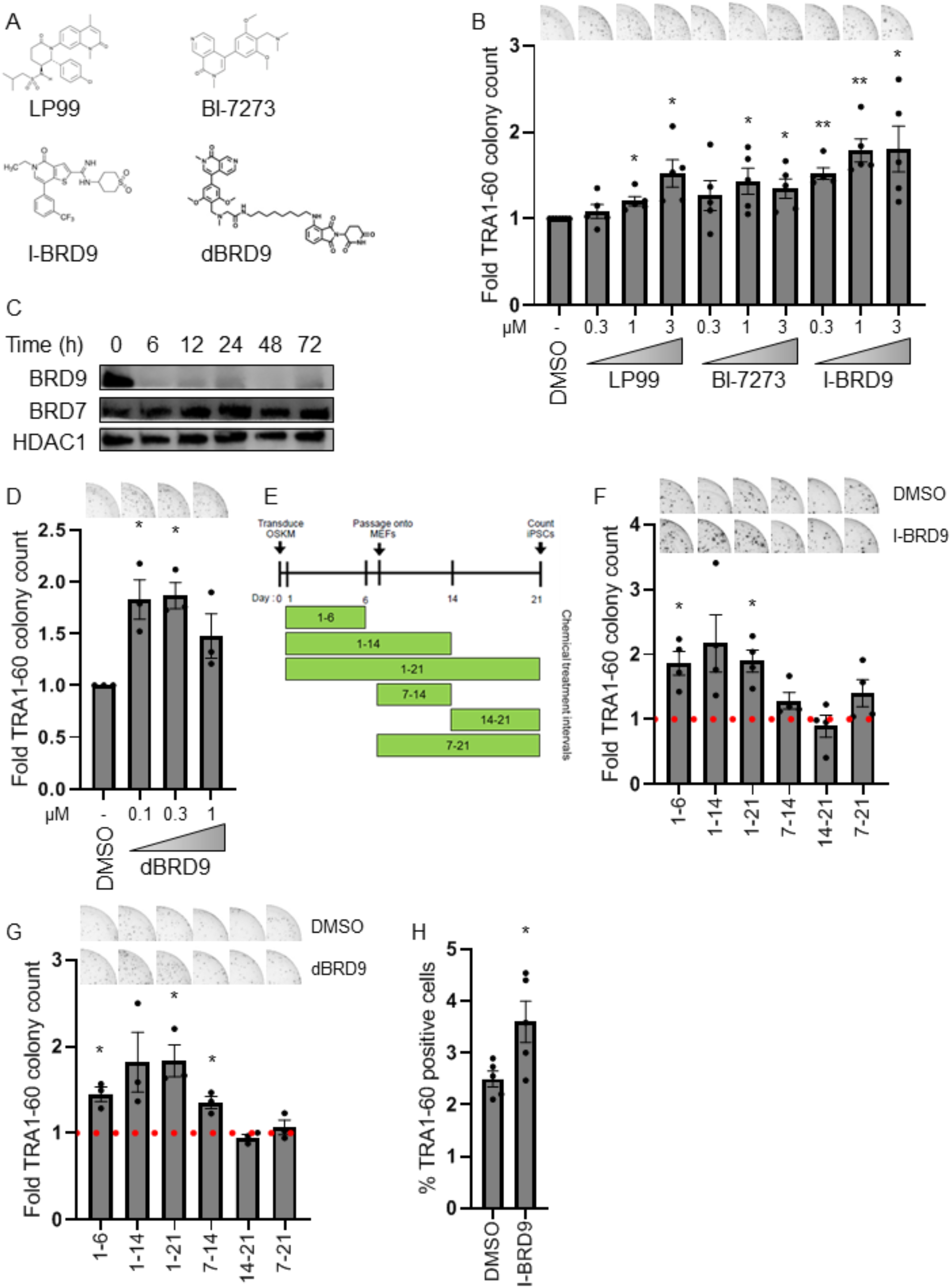
Bromodomain inhibition and degradation of BRD9 facilitate reprogramming. **(A)** Chemical structures of LP99, BI-7273, I-BRD9 and dBRD9. **(B)** Fold change in the number of TRA-1-60-positive colonies with the indicated compound treatments compared to DMSO control. Representative well images are shown above. Bar graphs show the mean and error bars represent standard error of mean. n=5, biological replicates for each treatment with 3 technical replicates. p-values are calculated by one sample t-test for mu=1. * denotes p<0.05, ** denotes p<0.005, exact p-values from left to right are: 0.3555, 0.0175, 0.0294, 0.1954, 0.0443, 0.0364, 0.0015, 0.0042, 0.0387. **(C)** Western blots for BRD9 and BRD7 at indicated time points after treatment with 0.3 µM dBRD9. HDAC1 serves as loading control. **(D)** Fold change in the number of TRA-1-60-positive colonies with increasing concentrations of dBRD9 compared to DMSO. Representative well images are above the graph. Bar graphs show the mean and error bars represent standard error of mean. n=3, biological replicates. p-values are calculated by one sample t-test for mu=1. * denotes p<0.05, exact p-values from left to right are: 0.0488, 0.0204, 0.1550. **(E)** Schematic depicting the time intervals for treatment of compounds during reprogramming. **(F)** Fold change in the number of TRA-1-60-positive colonies upon I-BRD9 treatment compared to DMSO between indicated days of reprogramming. Representative well images are above the graph. Bar graphs show the mean and error bars represent standard error of mean. n=4, independent biological replicates. p-values were calculated by one sample t-test for mu=1. * denotes p<0.05, exact p-values from left to right are: 0.0178, 0.0771, 0.0139, 0.1116, 0.5826, 0.1492. **(G)** Fold change in the number of TRA-1-60-positive colonies upon dBRD9 treatment compared to DMSO between indicated days of reprogramming. Representative well images are above the graph. Bar graphs show the mean and error bars represent standard error of mean. n=3, independent biological replicates. p-values are calculated by one sample t-test for mu=1. * denotes p<0.05, exact p-values from left to right are: 0.0332, 0.1417, 0.0456, 0.0376, 0.3099, 0.5208. **(H)** Percentage of TRA-1-60-positive cells on day 6 of reprogramming with DMSO and I-BRD9 treatment. Bar graphs show the mean and error bars represent standard error of mean. n=5, five biological replicates. p-values are calculated by two sample t-test. * denotes p<0.05, exact p-value: 0.0474.

Next, we took advantage of a recently described PROteolysis Targeting Chimera (PROTAC) targeting BRD9, dBRD9, to acutely deplete this protein (*31*). Time-course experiments showed that treatment with dBRD9 can dramatically decrease BRD9 protein levels for up to 72 hours without any effects on the closely related BRD7 (Fig. 2C). We observed that reprogramming efficiency increased up to 2-fold compared to control treatment even at the lowest concentration of 0.1 μM dBRD9 tested (Fig. 2D). A similar phenotype was observed in episomal plasmid-based reprogramming of an additional human dermal fibroblast line, indicating that BRD9 inhibition enhances iPSC generation independent of reprogramming strategy or cell line used (fig. S2E). Taken together, these results show that bromodomain inhibition or acute degradation of BRD9 increases human somatic cell reprogramming efficiency.

To understand how BRD9 inhibition promotes iPSC generation, we first determined when in the reprogramming process its inhibition has the maximal effect. A time-course treatment experiment with I-BRD9, dBRD9 and DMSO control was performed at different time-windows to evaluate which stage of reprogramming is responsive to BRD9 inhibition (Fig. 2E). Inhibition or degradation of BRD9 had the most effect on reprogramming efficiency when applied during the first 6 days after OSKM expression. As there were no further increases in the number of iPSCs with longer periods of chemical treatments, we concluded that BRD9 is a barrier for the initial stage of reprogramming (Fig. 2, F and G). In addition, the percentage of emerging TRA-1-60 positive cells on day 6 of reprogramming, as assessed by flow cytometry, was significantly higher in BRD9 bromodomain inhibitor-treated cultures compared to controls at this early time-point (Fig. 2H and fig. S2F). Taken together, these results indicate that the initial stage of reprogramming is most sensitive to BRD9 inhibition and that even a transient BRD9 inhibition is sufficient to increase the efficiency of human iPSC generation.

### BRD9 inhibition and degradation enable iPSC generation without KLF4 and c-MYC

We previously reported that inhibition of somatic barriers to reprogramming can enable human iPSC generation with fewer Yamanaka factors (*4, 10*). To investigate if BRD9 inhibition may have a similar effect, we carried out reprogramming with only OSK or OS. In both circumstances, reprogramming efficiency was increased with BRD9 inhibition (Fig. 3, A and B). PCR with vector-specific primers validated the absence of KLF4 and MYC transgenes in genomic DNAs of iPSCs derived from OS-transduced fibroblasts (Fig. 3C). Importantly, iPSCs derived by OS transduction could be stably propagated and exhibited pluripotency characteristics such as silencing of retroviral transgenes, expression pluripotency markers such as OCT4, SSEA4 and NANOG and ability to form teratomas containing differentiated cells from all three germ layers (Fig. 3, D, E and F). These results show that BRD9 inhibition can enable iPSC generation with fewer exogenous transcription factors.

**Figure 3.**
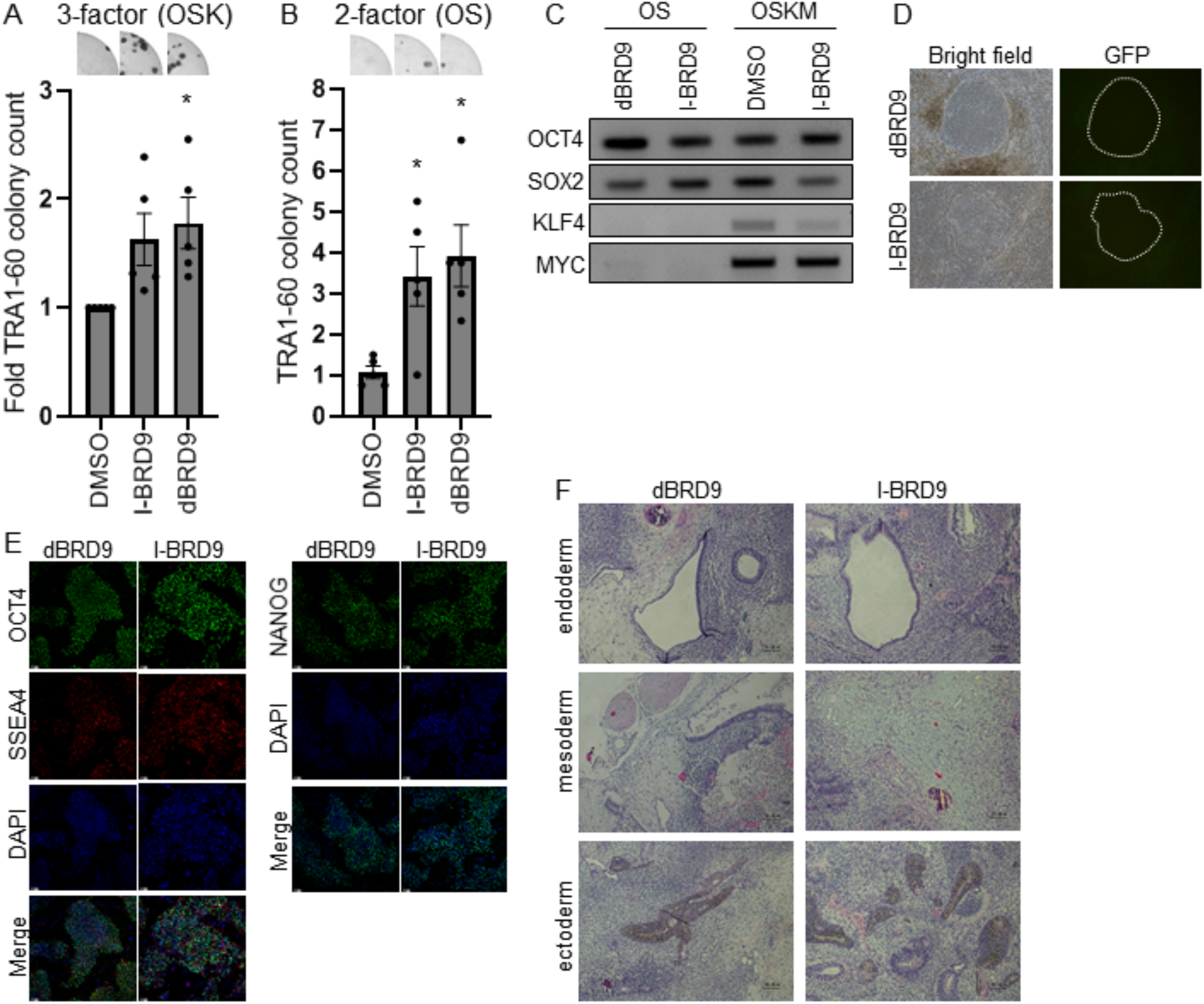
BRD9 inhibition and degradation enable iPSC generation without KLF4 and c-MYC. **(A)** Fold change in the number of TRA-1-60-positive colonies generated by OCT4, SOX2 and KLF4 (OSK) in the presence of DMSO, I-BRD9 or dBRD9. Representative well images are above the graph. Bar graphs show the mean and error bars represent standard error of mean. n=5, five biological replicates. p-values are calculated by one sample t-test for mu=1. Exact p-values from left to right are: 0.0583, 0.0297. **(B)** Number of TRA-1-60 positive colonies generated by OCT4 and SOX2 (OS) in the presence of DMSO, I-BRD9 or dBRD9. Representative well images are above the graph. n=5, biological replicates with at least 3 technical replicates. p-values are calculated by two sample t-test. * denotes p<0.05, exact p-values from left to right are: 0.0302, 0.0182. **(C)** Agarose gel images of PCR products for exogenous OCT4, SOX2, KLF4 and c-MYC transgenes in iPSCs generated from fibroblasts with indicated induction and treatments. **(D)** Phase contrast and GFP fluorescence images of colonies derived by OS induction and dBRD9 (upper) or I-BRD9 (lower) treatments showing typical iPSC morphology and silencing of retroviral GFP transgene. **(E)** OCT4, SSEA4 (left) and NANOG (right) immunofluorescence of iPSCs derived from OS expressing fibroblasts treated with indicated treatments. Hoechst 33324 was used to stain the nuclei. **(F)** Hematoxylin and eosin stained sections of teratomas of iPSCs derived from OS-expressing fibroblasts treated with dBRD9 or I-BRD9 show tissues from endoderm, ectoderm and mesoderm lineages.

### BRD9 is dispensable for human pluripotency induction and maintenance but required for mesendedorm differentiation

While small molecules allow for transient inhibition or degradation of BRD9 during reprogramming, sgRNA expression in fibroblasts may result in a permanent knockout in the resulting iPSCs. We therefore wished to determine whether the TRA-1-60-positive colonies generated from fibroblasts expressing *BRD9* sgRNAs have a complete absence of BRD9 protein. iPSC colonies derived from control and *BRD9* sgRNA expressing fibroblasts were expanded in culture and BRD9 protein levels were examined (Fig. 4A). 9 iPSC lines out of 14 generated from *BRD9* sgRNA expressing fibroblasts did not express BRD9 at all, suggesting a homozygous knockout. 4 iPSC lines had reduced protein expression suggestive of a heterozygous knockout and 1 iPSC colony retained wildtype levels of BRD9 (Fig. 4B). Complete knock-out clones robustly expressed OCT4, NANOG and SSEA4 similar to control iPSC lines (fig. S3A). The observation that the majority of the expanded clones expressed no BRD9 protein while exhibiting hallmarks of pluripotent stem cells suggests that loss of BRD9 is dispensable for human iPSC generation and propagation.

**Figure 4.**
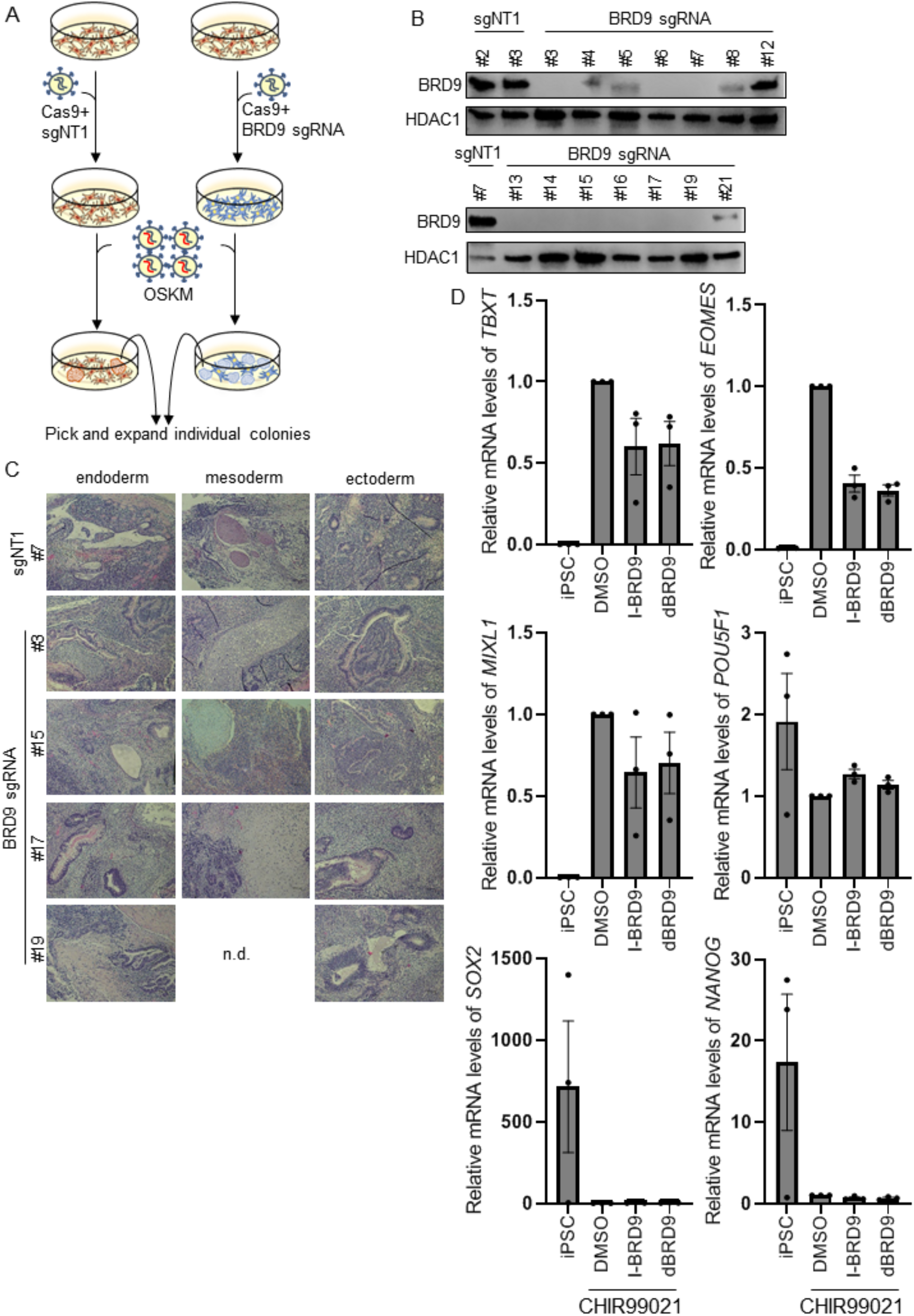
BRD9 is dispensable for human pluripotency induction and maintenance but required for mesendedorm differentiation. **(A)** Schematic for generation of BRD9 knockout and control iPSC lines **(B)** Western blot for BRD9 from iPSC clones generated from either sgNT1 or BRD9 sg823-expressing fibroblasts. HDAC1 serves as loading control. **(C)** Hematoxylin and eosin-stained sections of teratomas of iPSCs derived from sgNT1- or BRD9 sg823-expressing fibroblasts show tissues from endoderm, ectoderm and mesoderm lineages. n.d.: not detected. **(D)** RT-qPCR results for mRNA levels of *TBXT, EOMES, MIXL1, POU5F1, SOX2* and *NANOG* genes normalized to *ACTB* mRNA levels for indicated treatments. n=3, biological replicates. p-values are calculated by one sample t-test for mu=1. Exact p-values of I-BRD9 and dBRD9 treatments for *TBXT* expression are 0.1474 and 0.1069, respectively. Exact p-values of I-BRD9 and dBRD9 treatments for *EOMES* expression are 0.0076 and 0.0029, respectively. Exact p-values of I-BRD9 and dBRD9 treatments for *MIXL1* expression are 0.2429 and 0.2541, respectively. Exact p-values of I-BRD9 and dBRD9 treatments for *POU5F1* expression are 0.0436 and 0.1172, respectively. Exact p-values of I-BRD9 and dBRD9 treatments for *SOX2* expression are 0.2104 and 0.0590, respectively. Exact p-values of I-BRD9 and dBRD9 treatments for *NANOG* expression are 0.1376 and 0.0852, respectively.

To further characterize the pluripotency BRD9 knock-out hiPSCs, we investigated their differentiation capacity. BRD9-knockout iPSCs were able to form teratomas containing cells from all three germ layers (Fig. 4C). However, we observed that tissues from mesoderm lineage were less abundant in histological sections of teratomas generated by BRD9-knockout iPSCs. This observation led us to hypothesize that BRD9 may have a role in mesoderm differentiation. To test this, we differentiated two independent iPSC lines generated from two healthy donors into mesendoderm with WNT agonist CHIR99021 (*32*) and in the presence of small molecules targeting BRD9. Expression of *TBXT, EOMES* and *MIXL1*, well-established markers for mesendoderm, decreased upon BRD9 inhibition and degradation (Fig. 4D, fig. S3F). On the other hand, exit from pluripotency as judged by expression of *POU5F1, SOX2* and *NANOG* was not affected by BRD9 inhibition (Fig. 4D, fig. S3F). These results suggest that BRD9 is important for mesendodermal lineage commitment of human pluripotent stem cells.

Naïve mouse ESCs have been shown to be sensitive to BRD9 bromodomain inhibition when grown in serum and Lif (*27*), therefore we wished to further investigate the role of BRD9 in human pluripotent stem cells. We made use of an OCT4-GFP reporter human iPSC line (*33*) to monitor cell viability and self-renewal capacity upon BRD9 inhibition. BRD9 inhibitor or degrader treatment for 48 hours did not change the proliferation rate nor the percentage of OCT4-positive cells compared to controls (fig. S3, B, C and D). These results, in combination with the knockout iPSC lines, suggest that BRD9 is not required to maintain human pluripotency despite it being necessary for naïve pluripotency in the mouse (*27*). To specifically examine if the role of BRD9 in pluripotency acquisition is species-specific, we reprogrammed mouse embryonic fibroblasts in the presence of small molecules targeting BRD9. BRD9 inhibition and degradation did not increase murine somatic cell reprogramming; in fact, we observed a modest decrease in efficiency with the degrader (fig. S3E). These results show that BRD9, in contrast to its function in murine naive PSCs, is not required for the induction and maintenance of human pluripotency.

### BRD9 maintains somatic-specific gene expression and enhancer accessibility

Given that BRD9 inhibition is most effective in early reprogramming, where downregulation of the somatic cell gene expression program is a key rate-limiting step, we next sought to identify the transcriptional effects elicited by different modes of BRD9 inhibition. To this end, we performed mRNA-sequencing from fibroblasts treated with BI-7273, I-BRD9, dBRD9 as well as those expressing Cas9 and *BRD9* sgRNA. Small molecules targeting BRD9 differentially downregulated 928, 170, 577 genes (dBRD9, I-BRD9, BI-7273, respectively), of which 70 were common to all treatments (fig. S4A). Gene ontology (GO) analysis of this common set of genes revealed that they were highly enriched in cellular process linked to fibroblast identity and function, such as epithelial-to-mesenchymal transition (EMT), extracellular matrix components and adhesion (Fig. 5A). More broadly, Gene Set Enrichment Analysis (GSEA) indicated epithelial-to-mesenchymal transition (EMT) gene sets were among the top most negatively regulated gene sets upon I-BRD9 and dBRD9 treatments (Fig. 5B). This finding led us to ask specifically whether the fibroblast expression program as a whole was downregulated upon BRD9 inhibition. To test this notion, we performed GSEA using a fibroblast-related gene set (307 genes) that we generated based from gene expression profiles of human fibroblasts and their iPSC derivatives (*10*). All BRD9 perturbations resulted in a highly significant downregulation of the fibroblast-related gene set (Fig. 5C). In fact, the overall average expression values of genes in the fibroblast-related gene set were significantly downregulated upon BRD9 inhibition (Fig. 5D). In contrast, we did not observe positive enrichment of pluripotency-associated gene sets with any BRD9 perturbation, indicating that BRD9 inhibition on its own does not activate the pluripotency network (fig. S4B). Taken together, our transcriptomic analyses suggest that BRD9 acts as a barrier to reprogramming by sustaining starting cell type-specific gene expression.

**Figure 5.**
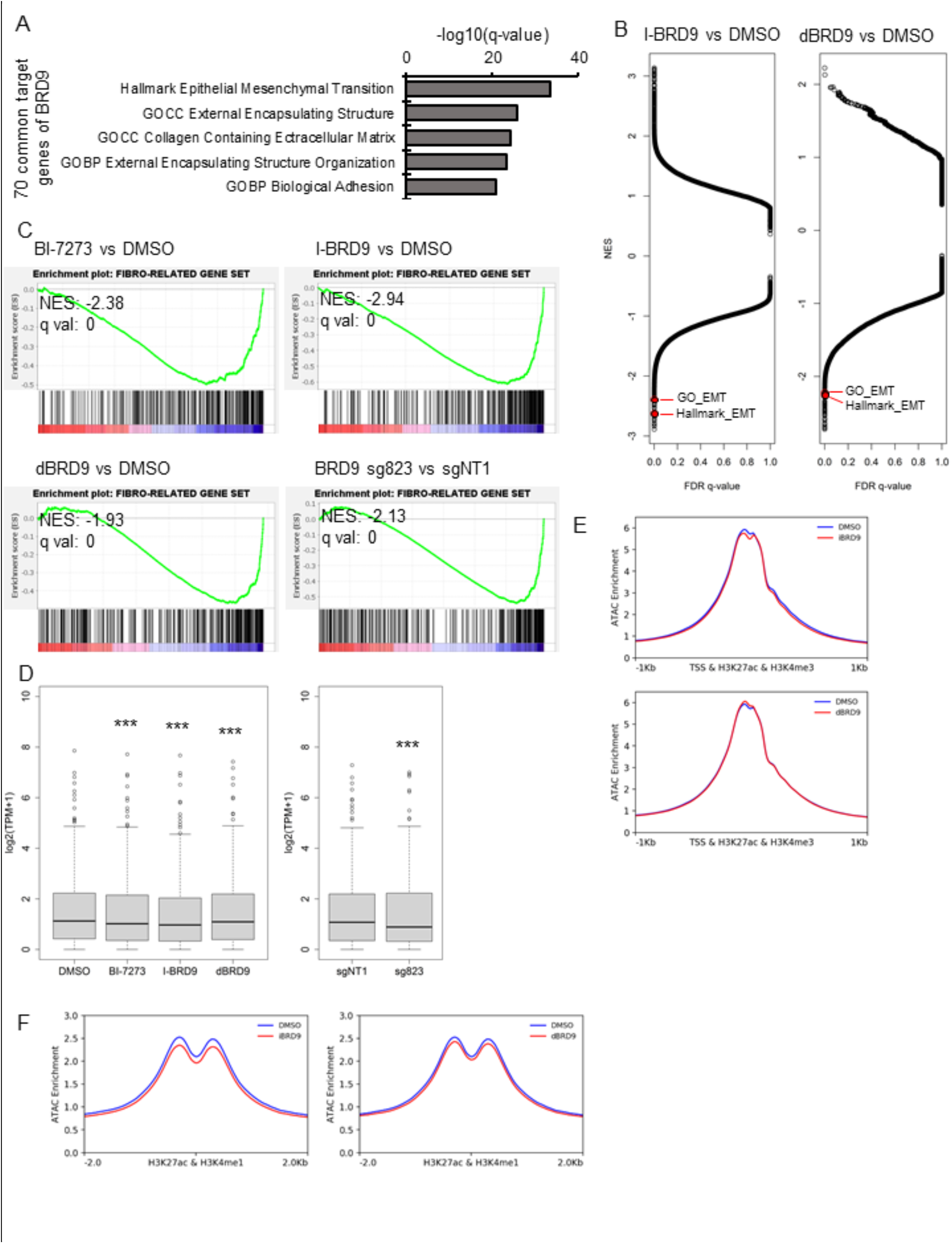
BRD9 maintains fibroblast-specific gene expression and enhancer accessibility. **(A)** Top five GO and Hallmark gene sets enriched in common downregulated genes upon dBRD9, BI-7273 and I-BRD9 treatments. Number of genes (n) in comparison were 70. p-values were calculated by hypergeometric distribution. **(B)** Gene set enrichment analysis on pre-ranked gene lists according to log2FC value for comparisons of fibroblasts treated with I-BRD9 and DMSO (left) and dBRD9 and DMSO (right) for all genesets available at The Molecular Signatures Database (MSigDB). Red circles indicate Hallmark_EMT and GO_EMT gene sets. **(C)** Gene set enrichment analysis (GSEA) of transcriptome data with indicated treatments for the fibroblast-related gene set. NES: normalized enrichment score, q val: False discovery rate (FDR) q-value. **(D)** Average expression levels of the 307 genes in fibroblast-related gene set across indicated fibroblasts. Whiskers indicate 95% confidence interval. p-values were calculated by Wilcoxon signed-rank test. *** denotes p<0.005, exact p-values from left to right are: 2.2e-16, 2.2e-16, 3.8e-10 and 5.057e-05. **(E)** Aggregate ATAC-seq plots from fibroblasts treated with I-BRD9 (top), dBRD9 (bottom) and DMSO on the +-1kb of transcription start site, H3K27ac and H3K4me3 summit. n=3, three biological replicates. **(F)** Aggregate ATAC-seq plots from fibroblasts treated with I-BRD9 (left), dBRD9 (right) and DMSO around +/-2kb of H3K27ac and H3K4me1 summit. n=3, three biological replicates.

To gain insight into how BRD9 functions to maintain expression of somatic-specific genes and test whether BRD9 has a role in maintaining chromatin accessibility at such loci, we performed ATAC-seq (Assay for Transposase-Accessible Chromatin using sequencing) in fibroblasts treated with I-BRD9 and dBRD9. Both inhibitors had no effect on accessible chromatin regions around promoters marked by overlap of H3K27ac and H3K4me3 (Fig. 5E). However, BRD9 inhibition or degradation reduced the accessibility of chromatin around putative active enhancers in fibroblasts as marked by overlap of H3K27ac with H3K4me1 (Fig. 5F). Importantly, such fibroblast-specific enhancers start to lose accessibility upon OSKM expression, suggesting BRD9 inhibition augments this process (fig. S4C) (*34*). These results indicate that BRD9 constitutes a barrier to reprogramming by maintaining accessibility of active enhancers in the starting cell populations.

### *MN1* and *ZBTB38* are BRD9 target genes that suppress reprogramming

Among the most consistently downregulated genes upon BRD9 inhibition were transcriptional regulators *MN1* and *ZBTB38* (Fig. 6A). MN1 has not been implicated in somatic cell reprogramming, but regulates palate development (*35*) and can act as co-factor for various transcription factors such as retinoic acid receptor/retinoic X receptor (RAR/RXR) (*36*). It is also implicated in transcriptional control of leukemic transformation in collaboration with DOT1L (*37*). ZBTB38 is predicted to be a master regulator in fibroblasts and is controlled by a fibroblast specific super-enhancer (*38*). We hypothesized that downregulation of these two factors soon after OSKM expression is necessary for efficient reprogramming. To test this notion, we overexpressed MN1 or ZBTB38 along with OSKM in human fibroblasts which resulted in a significant impairment in reprogramming efficiency (Fig. 6, B, C and D). Interestingly, *ZBTB38* expression in fibroblasts also significantly reduced the number of emerging TRA-1-60-positive cells on day 6 (Fig. 6E). Moreover, induction of pluripotency associated genes such as *NANOG, LEFTY2, LIN28A* were reduced at day 6 of reprogramming upon continued *ZBTB38* expression (Fig. 6F). These results BRD9 safeguards somatic cell identity and acts as a barrier to reprogramming in part by sustaining the expression these two transcriptional regulators (Fig 6G).

**Figure 6.**
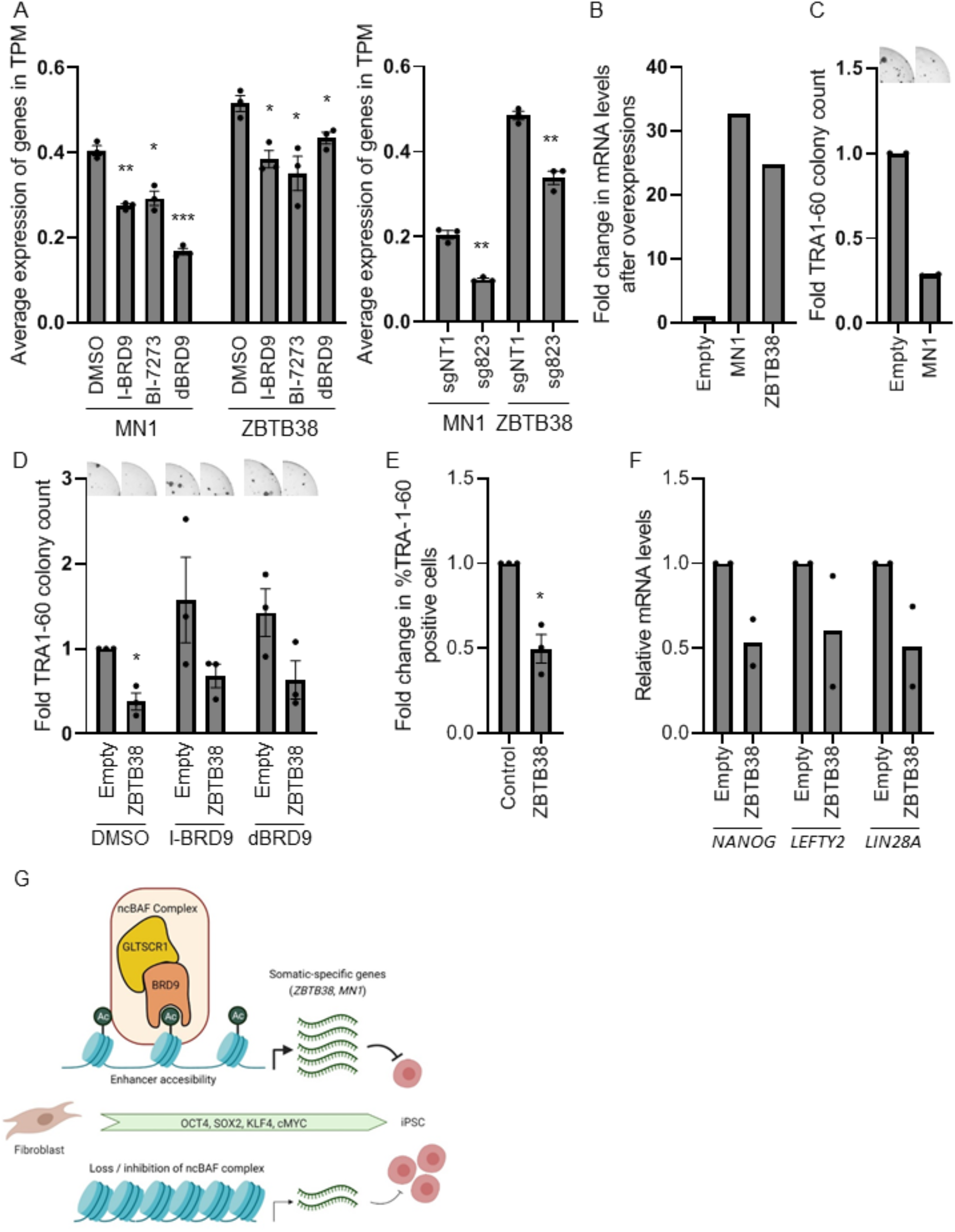
BRD9-regulated MN1 and ZBTB38 act as barrier to reprogramming. **(A)** Average expression of *MN1* and *ZBTB38* in TPM across indicated fibroblasts. n=3, biological replicates. p-values were calculated by two sample t-test. * denotes p<0.05, ** denotes p<0.005, *** denotes p<0.0005. Exact p-values for small molecule treatments compared to DMSO control from left to right are: 0.0031, 0.0076 and 0.0002 for *MN1* and 0.0089, 0.0378 and 0.0284 for *ZBTB38*. Exact p-value of BRD9 gRNA expressing cells to control cells is 0.0045 for *MN1* and 0.0026 for *ZBTB38*. **(B)** Fold change in relative *MN1 and ZBTB38* mRNA levels upon their overexpression in fibroblasts compared empty vector controls **(C)** Fold change in the number of TRA-1-60-positive colonies upon *MN1* overexpression. Representative well images are above the graph. Bar graphs show the mean. n=2, independent biological replicates. **(D)** Fold change in the number of TRA-1-60-positive colonies upon *ZBTB38* overexpression and indicated treatments. Representative well images are above the graph. Bar graphs show the mean and error bars represent standard error of mean. n=3, independent biological replicates. p-values are calculated by one sample t-test for mu=1. * denotes p<0.05, exact p-value is 0.0255. **(E)** Fold change in the percentage of TRA-1-60-positive cells on day 6 of reprogramming by comparing *ZBTB38* expressing cells to control. Bar graphs show the mean and error bars represent standard error of mean. n=3, three biological replicates. p-values were calculated by two sample t-test. * denotes p<0.05, exact p-value: 0.0272. **(F)** Fold change in relative mRNA levels for the *NANOG, LEFTY2* and *LIN28* genes on day 6 of reprogramming. Gene expression values were normalized to those observed in empty vector expressing fibroblasts. Bar graphs show the mean and dots indicate two biological replicates. **(G)** Model for BRD9-containing BAF complex’s role in maintaining somatic cell identity during reprogramming.

## Discussion

In this study, we investigated the role of BRD9 in human somatic cell reprogramming to pluripotency via a combination of genetic and chemical perturbation approaches. Knockdown of BRD9 via RNA interference or knockout via CRISPR/Cas9 increased human reprogramming efficiency. Importantly, the contrasting effects of BRD9 and BRD7 inhibition on reprograming efficiency indicate that the recently identified BRD9-containing ncBAF, but not BRD7-containing PBAF, is a major barrier to reprogramming. This notion is supported by our finding that loss of an additional specific member of ncBAF complex, GLTSCR1, has a similar positive effect on iPSC formation. To acutely block BRD9 function, we took advantage of selective bromodomain inhibitors and a PROTAC degrader, all of which significantly increased reprogramming efficiency and enabled iPSC generation in the absence of KLF4 and cMYC. Taken together, these findings demonstrate that BRD9-containing ncBAF complexes serve an important role in maintaining somatic cell identity.

Interestingly, we find that BRD9 is dispensable for induction and maintenance of human pluripotency. Fibroblasts expressing Cas9 and sgRNAs against *BRD9* could efficiently generate iPSCs that do not express any detectable BRD9 protein. Yet, such iPSC clones, and additional iPSCs treated with compounds targeting BRD9, could be propagated and expanded while retaining canonical properties of pluripotent stem cells. This is in contrast to the non-BRD9 containing ES-specific BAF complexes which are required for pluripotency and self-renewal (*39*). Interestingly, naïve mESCs have been shown to lose self-renewal capacity and enter into a primed, epiblast-like transcriptional state upon BRD9 inhibition (*27*). Human iPSCs are considered to be in a primed state (REF), our data indicate that BRD9 is dispensable for their maintenance. It is therefore likely that BAF complexes other than ncBAF such as non-BRD9 containing ES-specific complexes support human primed pluripotency. It is also worth noting that mouse and human reprogramming systems may have distinct species-specific features in part due to chromatin organization (*40, 41*). In fact, we observed that murine reprogramming was not enhanced upon BRD9 inhibition nor degradation. Our findings are consistent with a recent study in which I-BRD9 were found not to increase reprogramming efficiency from mouse embryonic fibroblasts (*42*). Collectively, these observations point to species-specific roles for BRD9 in somatic cell reprogramming and maintenance of pluripotency.

Genomic analyses indicate that BRD9 inhibition leads to broad downregulation of fibroblast-enriched genes which is accompanied by decreased chromatin accessibility across putative active enhancers and cell-type specific super-enhancers. These results align with previous studies which show BRD9 occupancy at distal enhancers (*26, 43*) and co-localization with CTCF (*23, 44*). In addition, BRD9 have recently been shown to be crucial for maintaining cell-type specific transcription programs in regulatory T-cells (*45*). We identified several key transcription factors such as *MN1* and *ZBTB38* as BRD9 targets. Our functional data, along with recent studies, establish these BRD9-regulated genes as important barriers to reprogramming (*46*).

The present findings add BRD9 bromodomain inhibitors and degraders to the arsenal of small molecule inhibitors that can be used to regulate and direct cell fate changes in human somatic cells. We also show that BRD9 inhibition can be combined with other modulators such as DOT1L inhibitors to boost reprogramming efficiency. Importantly, inhibition of DOT1L-mediated H3K79 methylation facilitates the generation of chemically induced pluripotent stem cells (ciPSCs) from mouse somatic cells (*47*) and result in a permissive epigenome state which enables reprogramming by alternative transcription factors (*48*). Identification of BRD9 as a safeguarding mechanism of cell identity suggests that combinatorial perturbations which include BRD9 inhibitors can enhance various reprogramming methods. Such approaches will likely lead to rapid silencing of the initial somatic program and potentially be used to derive human ciPSCs as well as direct lineage-converted cells with high efficiency.

## Materials and Methods

### Reprogramming assays

Human fibroblasts (dH1f) and reprogramming assays were performed as described previously (*10, 49*). Unless stated otherwise in the text, I-BRD9, LP99, BI-7273 and dBRD9 were used at final concentrations of 1 μM, 3 μM, 1 μM and 0.3 μM, respectively, for the first two weeks of reprogramming. EPZ-004777 was used at final concentration of 3 μM for the first week of reprogramming. HDF-A cells (ScienCell Research Laboratories, Catalog #2320) were reprogrammed with episomal vectors as described previously (*50*). For murine reprogramming, 25000 mouse embryonic fibroblasts (MEF) cells were seeded at each well of 12 well plate. The next day, they were transduced by lentiviruses containing a single stem cell cassette (*51*). Next day culture media was replenished with small molecules at the final concentrations indicated for human somatic cell reprogramming conditions. At 5th day of reprogramming, cells were transferred on inactivated MEFs. Next day and every other day until 14th day of reprogramming, culture media were replenished with mouse embryonic stem cell media (20% FBS, 1% NEAA, 1% Pen-Strep, 1% L-glutamine, 55nM 2-Mercaptoethanol, 1000 U/mL Leukemia Inhibitory Factor (LIF) (ESGRO, Catalog #ESG1107) in Knockout DMEM (Gibco, Catalog #10829-018)).

### iPSC culture and mesendodermal differentiation

Individual iPSC colonies generated from BRD9 sgRNA and Cas9 expressing fibroblasts were manually picked and cultured on inactivated MEFs with hESC media and ROCK inhibitor Y-27632 at a final concentration of 10 µM. After a few passages clones were transitioned to feeder-free conditions on matrigel (Corning, Catalog #354277)-coated plates with MEF-conditioned hESC media. iPSCs used for CHIR99021-induced mesendodermal differentiation experiments were picked from adult fibroblasts, electroporated with non-integrative episomal plasmids and cultured on feeder-free conditions on matrigel-coated plates with mTESR1 media (*50, 52*). When the cells reached 40-60% confluency, they were treated with DMSO, 1 µM I-BRD9 or 0.1 µM dBRD9 in 5 µM CHIR99021 in fibroblast media for 48 hours.

### Cloning

sgRNA and shRNA oligonucleotides (Table S1) were cloned into lentiCRISPR v2 (Addgene plasmid no. #52961) and into pSMP vector (Addgene plasmid no. #36394) respectively. shRNA cloning was performed as previously described with XhoI and EcoRI (*4*). In order to clone sgRNAs, lentiCRISPRv2 vector was digested with BsmBI enzyme and gel purified with MN PCR-Clean up Kit. Extracted DNA was treated with AP. Top and bottom guide RNAs were annealed with T4 Polynucleotide Kinase (3’ phosphatase minus, NEB) and T4 DNA Ligase Buffer (NEB) in a thermal cycler with the following settings: 37°C for 30 min, 95°C for 4 min and then ramp down to 25°C at 5°C/min. Annealed oligos were diluted to 1:200 and were used as inserts for ligating to digested lentiCRISPR v2 (50 ng) by T4 DNA Ligase (NEB) in 10X T4 DNA Ligase buffer (NEB). Ligation reaction were incubated for at least 2 hours at room temperature. Ligation mix was vortexed and spun down. 50 μl Stbl3 bacteria (NEB, Catalog: C737303) was mixed with 5 μl of the ligation reaction and left on ice for at least 15 minutes. Afterward, heat shock was applied at 42°C for 30 seconds in water bath. 150 μl LB was supplemented and cultured in 37°C at 225 rpm in a shaker for 1 hour. The transformed bacteria were spread on LB agar plate with ampicillin or carbenicillin.

### Virus production and transduction

Virus production was performed as described previously (*10*). Briefly, 2.5 × 10^6^ 293T cells per 10-cm dish were plated. The next day, cells were transfected with 2.5 µg viral vector, 0.25 µg pCMV-VSV-G (Addgene plasmid no. 8454), 2.25 µg psPAX2 (Addgene plasmid no. 12260) for lentivirus or pUMVC (Addgene plasmid no. 8449) for retroviruses using 20 µl FuGENE 6 (Promega) in 400 µl DMEM per plate. Supernatants were collected 48 h and 72 h post-transfection and filtered through 45-µm pore size filters. Human fibroblasts were doubly transduced with either shRNA or CRISPR vectors in consecutive days in the presence of 8 µg/ml protamine sulfate (Sigma). Transduced cells were selected by puromycin at 1 μg/ml.

### Western Blot

Cell pellets were resuspended in cytosolic lysis buffer (10mM HEPES pH 7.9, 10mM KCl, 0.1mM EDTA, 0.4% NP-40, Protease Inhibitor (1X) (Roche)) and shaken on ice for 15 minutes. After centrifugation at 3000g at 40C for 3 minutes, pellets were washed with cytosolic buffer again. Pellets were resuspended in nuclear lysis buffer (20mM HEPES pH 7.9, 0.4M NaCl, 1mM EDTA, 10% Glycerol, Protease Inhibitor (1X) (Roche)) and sonicated at amplitude 40 for 10 seconds twice. Supernatant containing nuclear fragments was collected after centrifugation at 15000g at 40C for 5 minutes. Lysates were incubated at 950C for 15 minutes with 4X laemmli buffer (BioRad) containing β-mercaptoethanol (BioRad). Boiled samples and protein marker (Bio-rad, Catalog: 161-0374) were loaded on gel (BioRad, Catalog: 456-1084).

Proteins were transferred on PVDF membrane (BioRad, Catalog: 1620177) by Bio-Rad tans-blot turbo transfer system at mixed weight transfer setting and incubated in 5% blotting grade blocker (BioRad Catalog: 1706404) solution for 1 hour. Membranes were incubated with BRD9 antibody (Active Motif, Catalog: 61537) at 1:1000 ratio, BRD7 antibody (Cell Signaling, Catalog: D9K2T) at 1:1000 ratio and HDAC1 antibody (Santa Cruz, Catalog: sc-7872) at 1:500 ratio overnight at 40C. Next day, membranes were washed with TBS-T for 15 minutes three times and incubated with secondary antibody (Abcam, ab97051) for 1 hour at room temperature. Membranes were washed with TBS-T for 15 minutes 3 times and incubated shortly with ECL western blotting substrate (ThermoFisher) before imaging at LI-COR FC.

### Genomic DNA PCR

Genomic DNA was extracted from cells with Nucleospin Tissue kit (catalog no. 740952.50) following manufacturer’s instructions. 200 ng of genomic DNA was amplified with DreamTaq DNA Polymerase (catalog no. K1081) and indicated primer pairs (Table S1). OCT4, SOX2, KLF4 and c-MYC transgene-containing amplicons were amplified with 30 cycles of 30 s at 95 °C, 30 s at 54.5 °C and 60 s at 72 °C. Amplicons containing GLTSCR1L sgRNA-induced in-dels were amplified using a BIO-RAD T100 thermocycler with 30 cycles of 30 s at 95 °C, 30 s at 66.7 °C and 60 s at 72 °C.

### T7 endonuclease assay

PCR products containing GLTSCR1L-induced in-dels were purified by MN PCR-Clean up Kit. 400 ng of purified DNA was denatured and heteroduplexed using NEBuffer 2 (catalog no. B7002S) in a total volume of 19 µL with the following settings in a thermocycler: 5 minutes at 95 °C, ramping down to 85 °C with 2 °C/s, ramping down to 25 °C with 0.1 °C/s. 1 µL T7 endonuclease (catalog no. M0302L) was added into the heteroduplexed DNA reaction and mixed thoroughly. The reaction was incubated at 37 °C for 1 hour and visualized in BIO-RAD Gel Doc XR+ after running on agarose gels.

### Flow cytometry

Cell surface TRA-1-60 expression was analyzed with an Accuri C6 flow cytometer (BD) using PE-conjugated anti-human TRA-1-60-R antibody (Biolegend, catalog no. 330610).

### Quantitative RT–PCR analyses

qPCR assays were performed as described previously (*10*) with the indicated primers (Table S1).

### Cell proliferation assay

OCT4-GFP iPSCs were seeded on matrigel-coated 96-well plates in mTESR1 media with ROCK inhibitor Y-27632 (10 µM). When cells reach 40-60% confluency, they were treated with DMSO, 1 µM I-BRD9 or 0.1 µM dBRD9 in mTESR1 media for 48 hours. Cell proliferation was measured with the CellTiter-Glo Luminescent Cell Viability Assay (Promega) according to manufacturer’s instructions using luminometric measurements with a plate reader (Synergy H1 Reader, BioTek).

### Immunostaining

TRA-1-60 staining was performed as previously described (*4*) and anti-SSEA1 antibody (Biolegend, catalog no. 125604) was used to quantify MEF reprogramming. For immunofluorescence-based characterization of iPSCs, cells were passaged onto mitomycin-C-treated MEFs in hESC medium and carried out as previously described (*10*). The antibodies used were OCT4 (Abcam, catalog no. ab19857), SSEA4/A647 (BD, catalog no. 560218) and NANOG (Abcam, catalog no. ab21624). For NANOG and OCT4, Alexa-488-conjugated secondary antibodies (Molecular Probes) were used. Nuclei were stained with VECTASHIELD Antifade mounting medium with DAPI (H-1200-10).

### Teratoma assays

All experiments were carried out under a protocol approved by Koç University Animal Experiments Ethics Committee and all relevant ethical regulations were complied with. Teratoma injections were performed as previously described (*53*).

### RNA-sequencing and analysis

Fibroblasts were treated for 5 days with DMSO, BI-7273 (1 μM), I-BRD9 (1 μM) and dBRD9 (0.3 μM). In parallel, control gRNA (gNT1) and BRD9 g823-expressing cells were used. Total RNA was prepared using Direct-zol kit according to manufacturer’s instructions (Zymo Research). Libraries were prepared using NEBNext Poly(A) mRNA Magnetic Isolation Module from the NEBNext ultra-directional RNA kit to create a first stranded library. Reads were mapped to hg19 built-in genome by HISAT2 after assessing their quality by FastQC. DESeq2 package was used to find differentially expressed genes between samples. Genes were considered as differentially regulated based on adjusted p-value<0.05. The formation of fibroblast-related gene set was described previously (*10*). Rank-ordered gene lists were used for gene-set enrichment analysis (*54*).

### ATAC-Sequencing and analysis

ATAC-seq was performed using 100,000 cells for the transposition reaction as described previously (*55*) using in-house-produced Tn5 transposase. Subsequently, samples were purified using the GeneJET PCR purification kit (Thermo). PCR amplification was performed using the following protocol: 3 min at 72 °C, 30 s at 98 °C and 11 cycles of 10 s at 98 °C, 30 s at 63 °C, and 3 min at 72 °C. The samples were then purified using the GeneJET PCR purification kit and eluted with 20 μl of TE buffer. Samples were then run on a Tapestation (Agilent) to determine library size and quantification before paired-end (2 × 41 bp) sequencing on a NextSeq 500 (Illumina) platform. Sequence reads were quality controlled with FastQC and mapped to the most recent human reference genome (hg38) using bowtie (*56*), allowing a maximum of 2 mismatches and only uniquely mapped reads. Reads with mapping quality score below 30 and blacklisted regions were filtered. DeepTools (*57*) ComputeMatrix and plotProfile commands were used to generate aggregate ATAC plots. For this, BigWig files were generated using deepTools bamCoverage command, eliminating duplicates and normalizing by sequencing depth and effective genome size. Intersecting regions among fibroblast ChIP peaks for H3K27ac, H3K4me1 and H3K4me3 were identified using previously published fibroblast ChIP-seq data (*10*), calling the peaks using macs2 (*58*) with the parameters - -keep-dup 1 and –broad, and finding the intersecting regions using bedtools (*59*) intersect.

### Statistical Analysis

Statistical analysis was performed using GraphPad Prism 8 for t-tests or R 4.0.2 for Wilcoxon-signed rank test. The test details, number of biological replicates and exact p-values were provided in related figure legends.

## Acknowledgments

We would like to thank Ahmet Kocabay and Ali Cihan Taşkın for help with mouse experiments, Arzu Ruacan (Koç University, School of Medicine, Department of Pathology) for examination of histological sections. The authors gratefully acknowledge use of the services and facilities of the Koç University Research Center for Translational Medicine (KUTTAM), funded by the Republic of Turkey Ministry of Development. The content is solely the responsibility of the authors and does not necessarily represent the official views of the Ministry of Development. KS and DHA were supported by TUBITAK BIDEB Scholarship.

## Funding

Newton Advanced Fellowship (TTO)

TUBITAK Projects 231S182 and 219Z209 (TTO)

Arthritis Research UK, program grant 20522 (UO) Cancer Research UK (UO)

LEAN project of the Leducq Foundation (UO)

People Programme Marie Curie Actions of the European Union’s Seventh Framework

Programme FP7/2007-2013 under REA grant agreement n° 609305 (UO)

## Author contributions

Conceptualization: KS, GGS, UO, TTO

Software: KS, ADC, APC

Formal analysis: KS, GGS, ADC, TTO

Investigation: KS, GGS, MP, SK, HC, ESA, HAD

Resources: APC, JED, ABD, LHS, JQ

Data Curation: KS, ADC, MP, APC

Writing—original draft: KS, TTO

Writing—review & editing: GGS, ADC, SK, JQ, UO

Supervision: JQ, UO, TTO

Funding acquisition: UO, TTO

## Data and materials availability

RNA-sequencing and ATAC-sequencing data are deposited to the NCBI GEO database with the accession number GSE161640. Necessary information to produce materials and replicate analysis is provided in the main text or the supplementary materials.

